# Nothing but lies: improving the validity of neural predictors of deception

**DOI:** 10.1101/2024.05.08.593230

**Authors:** Sangil Lee, Runxuan Niu, Lusha Zhu, Andrew Kayser, Ming Hsu

## Abstract

Deception is a universal human behavior. Yet longstanding skepticism about the validity of measures used to understand the biological mechanisms underlying deceptive behavior has relegated such studies to the scientific periphery. Here we address these fundamental questions by applying novel machine learning methods and functional neuroimaging to signaling games capturing motivated deception in human participants. First, we develop an approach to test for the presence of confounding processes and thereby validate past skepticism by showing that much of the predictive power of neural predictors trained on deception data comes from confounding processes. Second, we show that the presence of confounding signals need not be fatal, and we improve the validity of our neural predictor via a novel machine learning procedure that identifies and removes these confounding signals. Together, these findings point to a scientific approach for studying a neglected class of behavior, with important methodological and societal implications.

## Introduction

Claims that scientifically-based methods can detect deception have been made since at least the early part of the 20th century. Yet from the start, such proposals, typically driven by forensic goals, have been met with intense skepticism from the scientific community, in large part due to uncertainty about the nature of the measured signals (*1–5*). Indeed, a conundrum dating to the earliest attempts at detecting deception involves how to rule out the myriad alternative processes, such as those involved in arousal, weighing of risks and rewards, and belief inference, that often co-occur with deception but are not necessarily themselves indicative of deception (*4–8*).

Scientifically, this continued lack of progress in our ability to assess and exclude validity threats has severely impeded progress in understanding the neural bases of deceptive and honest behaviors, which are central in studies of mate selection, social communication, and economic exchange, among others (*9–14*). As described by the National Research Council, “An indication of the state of the field is the fact that the validity questions that scientists raise today include many of the same ones that were first articulated in criticisms of Marston’s original work in 1917.” (*5*)

In recent years, however, there has been renewed excitement in the possibility of a more scientifically-grounded understanding of the neural basis of deception. Due to a confluence of advances in behavioral, neural, and statistical methods, new analysis approaches offer formal, testable ways to decode mental states from brain data (*15–17*), and in particular the application of machine learning pattern analysis techniques to economic signaling games (*18–20*). Such games have now been extensively studied in the economic and biological sciences in order to model goal-directed communication between agents, including motivated deception (*9–12*).

Here we seek to build upon these advances by developing a set of novel methodological and statistical tools to enable researchers to systematically test and improve the validity of putative predictors of deception. As a first step toward establishing criterion validity, we tested the accuracy of a neural predictor by applying multivariate decoding methods to functional neuroimaging data while participants made decisions about whether to send honest or deceptive messages to another participant. Our initial results show that, once trained on behavior in this game, a whole-brain neural predictor is capable of distinguishing between deceptive and honest behavior at rates significantly higher than chance.

Second, and perhaps more importantly, we set out to address questions about the construct validity of our neural predictor – i.e. how and why our predictor works. Whereas accuracy metrics ask whether our neural predictor can make accurate predictions, construct validity asks whether our neural predictor is truly measuring the underlying process it purports to measure. Despite the acknowledged importance of construct validity, and the many methods of detection proposed over the past 100 years, there has been surprisingly little effort in attempting to assess construct validity of these methods, nor even an agreement on what constitutes scientific evidence for or against the presence of validity threats (*4*, *5*, *21*).

In particular, we focus on the aspect of construct validity that pertains to discriminant validity (*22*, *23*), which assesses the extent to which our deception predictor is driven by processes not specific to deception. For this reason, we introduced a second, isomorphic, signaling game in which players can achieve the same ends (i.e., payoffs) as in the deception game, but via non-deceptive means. Critically, if the putative deception predictor “overgeneralizes” – meaning that its predictions are also correlated with signals underlying this control game – this result would provide strong evidence that its predictive power is significantly driven by processes held in common between the two games – e,g., self-interested motives, belief inference, arousal associated with violation of social norms, or others – rather than those specific to deception itself (*17*).

Using the neural predictor trained on the deception game, we show that its construct validity is severely compromised by the fact that this predictor also predicts “merely selfish” (i.e., selfish but not deceptive) behavior in the control game. Indeed, the magnitude of this effect is such that the prediction rate in the control task is statistically indistinguishable from that for the task of interest. Moreover, performance falls to chance when the predictor is asked to distinguish between (a) deceptive choices and (b) selfish but not deceptive choices. Lastly, at the neural level, we find that overgeneralization is pervasive across the brain, such that many regions that predict deception are at least partially driven by signals common to the control task.

Having identified the presence of confounding processes, we further investigated the extent to which the influence of any identified confounding processes can be removed or mitigated. This step is particularly important because the presence of confounding signals need not rule out the possible presence of a coexisting deception-specific signal. If so, we may be able to improve the validity of the neural predictor by purging the set of signals common to both tasks. To this end, we develop a novel statistical approach where, in addition to the typical goal of predicting the behavior of interest, the predictor also incorporates a second goal enforcing chance performance in the control task.

We show that compared to alternative methods, this “dual-goal tuning” approach is able to construct a whole-brain deception predictor that predicts deception, but does not rely on neural patterns held in common with the control task, rendering it capable of distinguishing between deceptive and merely selfish behavior. Additionally, this method uncovers substantial variation in the extent to which deception-specific signals can be recovered: whereas dual-goal tuning in some regions, such as the primary visual cortex, entirely purges presumptive predictive signals – suggesting that predictive accuracy in these regions is driven by processes other than deception – other regions, including ones previously implicated in meta-analyses of deception (such as the dorsal anterior cingulate cortex and superior frontal gyrus), retain significant predictive power after correction. Together, these findings potentially enhance the scientific rigor of studies that assess the neural basis of deceptive and honest behaviors, and they represent an important step forward in building a firm scientific foundation necessary for ongoing progress in detecting deception in forensic settings (*1–7*).

## Results

### Signaling game approach to dissociate processes underlying deception

To identify a set of processes that co-occur with but are not necessarily indicative of deception, we used a signaling game approach that has been extensively employed in behavioral economics and evolutionary biology to capture the role of communication in economic interactions, including tradeoffs between honesty and deception (*9*, *11–13*, *24*). Specifically, we designed a pair of isomorphic signaling games that differed only in the extent to which players’ actions could be assigned a truth value (*11–13*). In both games, the participant (the sender) is presented with two potential allocations of monetary gains for themselves and a counterpart (the receiver). On each trial, one allocation provides a larger payoff to the sender, while the other provides a larger payoff to the receiver. Critically, in the deception task, senders must choose between two messages that are verifiably truthful or false (e.g., “Option A will earn you more money than B”; **Figure 1A** - deception task). In contrast, in the control task, the senders’ messages do not have a truth value (e.g., “I prefer that you choose option A”; **Figure 1A** - control task).

**Figure 1.**
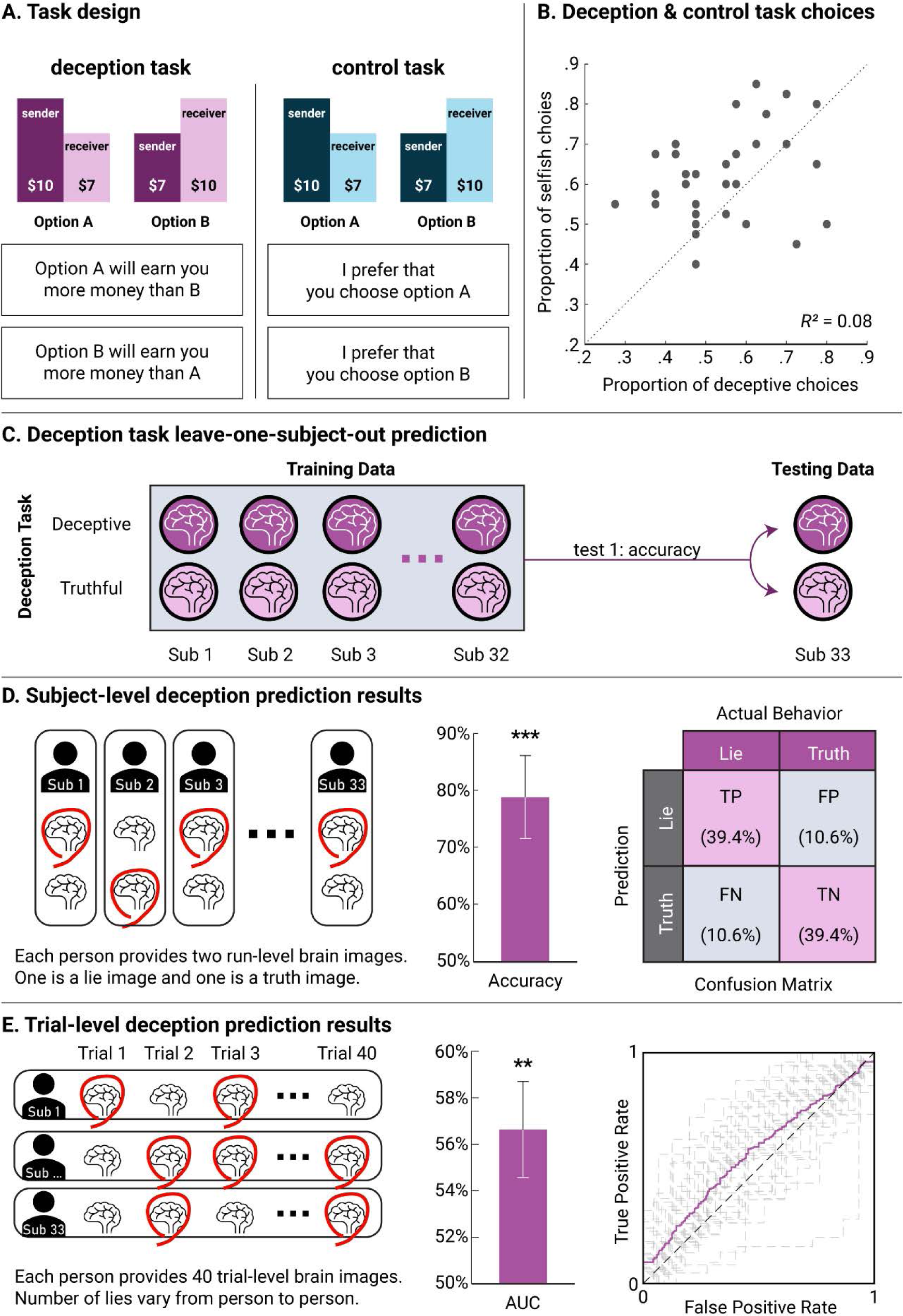
(**A**) Experimental task design dissociating deception from basic social decision-making processes. Both task conditions (deception and control) involve allocations of monetary gains between the participant and a counterpart. Participants always played the role of the message sender, who sends one of two messages to the receiver. The deception task requires senders choose between two messages that are verifiably truthful or false (e.g., “Option A will earn you more money than B”). In the Control Task, the senders’ messages do not have a truth value (e.g., “I prefer that you choose option A”). Importantly, participants are informed that as the sender, they can see the options but cannot make the choice between them, while the receiver cannot see the options but is responsible for the choice. This manipulation thus renders the receiver completely reliant upon the sender for any information about the choice (see Supplement for further task details). (**B**) Scatterplot of the proportion of deceptive choices from the main task and the proportion of selfish choices in the control task. That behavior in the deception task could be partially dissociated from those of the control task suggest contribution of processes that are not simply reducible to processes captured in the control task. (**C**) Schematic of test 1: leave-one-out cross-validation procedure for within-task prediction. (**D**) Predictors were trained on subjects’ run-level average neural activity for each choice type (two images per task for each subject). Neural predictors of deception predicted deceptive behavior at rates significantly greater than chance (78.8%±7.24% (mean±SE), p<0.001). (**E**) In trial-level prediction, each trial’s activity was estimated separately and predictions were made at the trial level. The neural predictor of trial-level deception also showed significant prediction of deceptive choices (AUC = 56.6%±2.1% (mean±SE), p = 0.004). ** *p* < .01, *** *p* < .001.

This design incorporates two important features that together help to identify deception-specific processes. First, in contrast with previous paradigms involving instructed lies (*25–29*), behavior in signaling games captures the idea that honesty and deception are properties of the communicative signals that agents send to one another in the service of some economic or evolutionary value. Second, and more importantly, the inclusion of an isomorphic control game allows us to identify the set of processes that are also present in other non-deceptive decisions—for example, those associated with weighing costs and benefits to oneself, or concerns for fairness—and thus that are not specific to deception per se.

Consistent with previous findings showing that processes underlying deception can be dissociated from those underlying other types of norm violations (*11–13*), there was a significant difference in how participants behaved in the two conditions. In particular, the need to send a deceptive message reduced the proportion of messages recommending the selfish option to the receiver (54.6%) compared to the control condition (61.9%; paired t-test *p* = .0074). Notably, individual differences in sensitivity to social norms around self-favoring and honest responses could be dissociated across the two tasks, indicating that the message manipulation differentially impacted behavior: those participants who made more deceptive choices in the deception task were not necessarily the ones who made more selfish choices in the control task (**Figure 1B**; *r* = 0.28, *p* = 0.11).

### Neural predictor distinguishing between deceptive and honest behavior

Next, using functional neuroimaging data, we sought to assess the extent to which a whole-brain neural predictor, once trained on neural responses associated with deceptive and honest behavior (**Figure 1C**), could predict deception in holdout data using brain activity alone (*18*). Specifically, training and testing were performed at both subject- and trial-levels. In subject-level prediction, we estimated, for each subject, one image associated with truthful behavior and one with deceptive behavior, by estimating average brain activity across the same trial types (*30*, *31*). We found that the neural predictor was able to correctly distinguish the two images at rates significantly greater than chance (78.8%, *p* < .001; **Figure 1D**). In trial-level prediction, separate images were estimated for each choice (*32–34*). Using the area under the receiver operating characteristic curve (AUC) as a measure of overall performance of the classifier across all possible thresholds, where 50% corresponds to chance performance and 100% corresponds to perfect classification, we found that the neural predictor performed significantly better than chance at the trial level (average AUC = 56.6%, p = 0.004; **Figure 1E**).

### Significant presence of confounding signals

Although this neural predictor of deception was able to show significant discrimination between deceptive and honest behavior, it is possible that at least some of the predictive signal is not related to deception, but rather to confounding processes. We sought to test for this possibility by building upon recent methods that ask whether a neural predictor of a particular cognitive state shares signals with other processes (*17*, *34*, *35*). For example, researchers might develop a neural predictor in one dataset, then apply that predictor to another dataset that utilizes a different task in order to (putatively) assess the same underlying construct. A significant correlation between the predictive signal and the behavior of the new task (i.e., generalization) argues that the predictor of one construct incorporates the other construct, whereas a lack of generalization suggests independence of the constructs in question.

By testing the extent to which the neural predictor of deception trained on the deception task can also distinguish between behavior in the isomorphic control signaling game that does not involve deception, this approach provides one way to identify threats to discriminant validity in the neural predictor (**Figure 2A**) – in other words, by evaluating whether measures that should not be correlated are indeed uncorrelated (*36*). In contrast, if a neural predictor trained on the deception game produces signals that are also associated with non-deceptive behaviors, we can conclude that this “naïve” predictor is at least partially driven by processes held in common between the two games – e,g., self-interested motives, belief inference, arousal associated with violation of social norms, or others – rather than those specific to deception itself.

**Figure 2.**
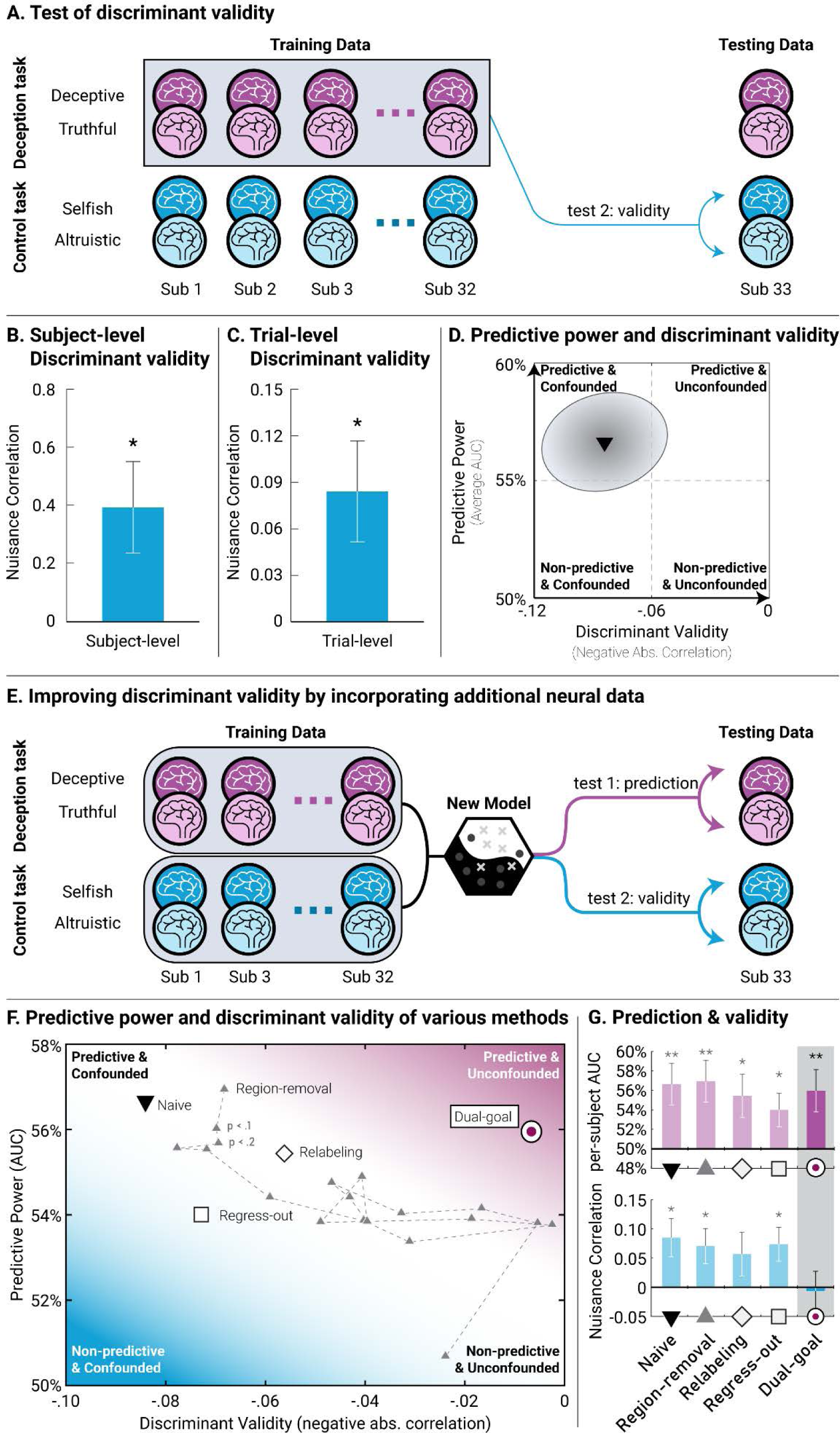
(**A**) Schematic of test 2. Discriminant validity is tested by using the control task as testing data, rather than deception task trials as in test 1. A predictor that is associated with behavior in the control task indicates a lack of discriminant validity. (**B**) In subject-level testing, the neural predictor of deception showed a lack of discriminant validity, as it was significantly correlated with behavior in the control game (mean r = 0.39, *p* = 0.021). (**C**) Neural predictors showed a lack of discriminant validity in trial-level prediction as well (mean r = 0.084, p=0.014). (**D**) Combining test 1 and test 2, an ideal neural predictor should be located in the upper right quadrant where it is predictive of deception (ordinate) but uncorrelated with other behaviors (abscissa). In contrast, the neural predictor trained on deception data was located in the upper left quadrant – i.e. it was able to demonstrate predictive accuracy when tested on deception and honest trials, but underlying signals contained processes held in common with the control task. (**E**) Constructing and comparing methods that seek to improve discriminant validity by incorporating the control task into training data. (**F**) As in (D), predictive accuracy versus discriminant validity, compared across different neural predictors. Generally, all methods used, when compared with the naive predictor, showed some tradeoff between predictive power and discriminant validity, with dual-goal tuning providing the most favorable tradeoff in terms of power loss and discriminant validity. (**G**) Bar graphs of prediction performances and validity tests for methods shown in panel **F**. * *p* < .05, ** *p* < .01, *** *p* < .001.

We found strong evidence that the deception predictor incorporates signals that are shared with the control task. Specifically, the neural signal produced by applying a deception predictor to a control task was positively correlated with selfish but non-deceptive choices in the control task (subject-level: r = 0.39, p = 0.021, **Figure 2B**; trial-level r = 0.084, p = 0.014, **Figure 2C**). Thus, despite the significant predictive accuracy in the deception task, this overgeneralization provides strong evidence of a lack of discriminant validity for this predictor (**Figure 2D**). Indeed, the strength of this overgeneralization is such that if one were to use the putative deception predictor to classify the choices in the control task, the prediction accuracy would be statistically indistinguishable from its performance in the deception task at both the subject prediction level (78.8% vs. 69.7%; *p* = 0.45) and the single-trial prediction level (56.6% vs. 55.4%; *p* = 0.71).

### Methods to control for confounding processes

While the presence of overgeneralization shows the vulnerability of our neural predictor to confounding, it also offers an opportunity to improve its validity by identifying the set of such signals that can be removed if handled appropriately. To explore this possibility, we considered four potential solutions by incorporating data from the control task into the training process (**Figure 2E**).

First, if the loci containing confounded signals are spatially separate from those carrying signals of interest, a ‘region removal’ method can be used to improve the validity of the neural predictor by removing regions that carry strong confounding signals. Second, we considered a data ‘relabeling’ method, also known as the binary relevance method (*37*), that is often used in the machine-learning literature in the context of multi-class prediction problems. In our setting, this method works by assigning all trials of the control task to be ‘truth’ trials, with the only ‘lie’ trials supplied by the main task. By explicitly defining both control categories as ‘truth’ trials, this method ensures that the predictor is trained to down-weight any confounding signals present in the control task categories (i.e., selfish vs. altruistic). Third, we considered a ‘regress-out’ method, which attempts to focus the neural predictor on signal variation unique to the deception task. This method takes advantage of the isomorphic nature of our task, in that both tasks have the same number of trials and choices in the two tasks can be paired. As such, we can regress behavioral variation in the control task out of the deception task. (However, the trial pairing requirement limits this approach to trial-level prediction; see **supplemental materials** for a more in-depth discussion of the application of alternative regress-out methods.)

Finally, we developed a fourth method that seeks to directly control for cross-task (over)generalization. Unlike the other methods considered, but akin to post-training procedures developed in machine learning models to ensure fairness or reduce bias (*38*), this approach incorporates a second objective that explicitly seeks to remove any information that is predictive of the control task. Formally, given two datasets, (***X****_1_, **Y**_1_*) and (***X****_2_*, ***Y****_2_*), a deception-only predictor would be consistent with a linear model ***b*** such that ***X****_1_**b*** is predictive of ***Y****_1_*, but ***X****_2_**b*** is not correlated with ***Y****_2_*. If both ***X*** and ***Y*** have been mean-centered, the latter condition can be expressed as follows:

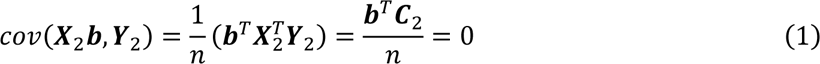

Hence cross-task generalization is caused by the similarity (i.e., the inner product) between the prediction map ***b*** and the covariance map ***C****_2_* of the nuisance (control) task. We therefore develop a two-step approach in which the predictor ***b*** is constructed first, then orthogonalized with regards to the nuisance covariance map using a Gram-Schmidt procedure. This method instantiates a ‘dual-goal tuning’ approach in which the hyperparameters for constructing the initial predictor ***b*** are chosen to maximize predictive performance in dataset 1, while a second hyperparameter is chosen to set cross-task generalization performance in dataset 2 to zero. In cost function-based methods such as ridge regression, one can reformulate this approach in the form of a penalty term for cross-task generalization (see **supplemental materials**).

### Improving neural predictor discriminant validity

Ideally, a successful method should remove any correlation with non-deceptive behaviors while retaining as much of its predictive power in the task of interest as possible—i.e., shifting the naïve predictor away from the “predictive confounded” quadrant and toward the “predictive unconfounded” region (**Figure 2D**). Using a cutoff p-value of 0.05 for the region-removal method, where voxels significantly correlated with the control task are masked out, we found negligible change in performance as compared to the naïve approach (**Figure 2F & G**). A more systematic examination of different cutoff values (p < .1, .2, …, .99) further showed that, while confounding signals were indeed removed as more regions were masked out, this improvement was largely achieved at the expense of accuracy in the task of interest (**Figure 2F**). Similarly, the relabeling approach and the regress-out methods were only able to remove confounding signals at the expense of reduced accuracy (**Figure 2F & G**).

In contrast to the poor to mixed performance of the previous three approaches, we found that the dual-goal tuning approach significantly reduced overgeneralization compared to the naive method (*p* < .001) and was in fact nearly able to eliminate it completely (*r* = −0.0067, *p* = 0.85); while retaining predictive power for deceptive choices (single-trial prediction: 56.0%, *p* = 0.01; **Figure 2F & G**). This difference reflects an important distinction between the other approaches and dual-goal tuning, in that the latter required much less tradeoff between predictive performance in the two tasks.

### Distinguishing between deceptive and selfish behavior

A potentially important limitation of inferring discriminant validity based on removal or absence of overgeneralization is that it relies on accepting the null hypothesis. Such conclusions are known to be problematic, as the null may fail to be rejected simply because the study lacked statistical power. To address this possibility, we constructed a positive test involving a “high confound” testing set consisting of deceptive and selfish trials.

Here, a helpful analogy can be made with pregnancy tests, which when used in the general population are known to be highly sensitive and specific. However, because the test does not measure pregnancy per se, but levels of β-hCG hormone, the test is known to perform poorly if used in a ‘high-confound’ testing set in which pregnant women are intermixed with populations with syndromes causing abnormally high levels of β-hCG, such as trophoblastic disease and certain cancers (*39*, *40*). Thus, just as a pregnancy test with improved discriminant validity can be demonstrated by successfully distinguishing between pregnant and other individuals with elevated β-hCG levels (*39*, *40*), we can provide positive evidence that discriminant validity has improved by showing that the corrected neural predictor can distinguish between deceptive and merely selfish behavior (**Figure 3A**).

**Figure 3.**
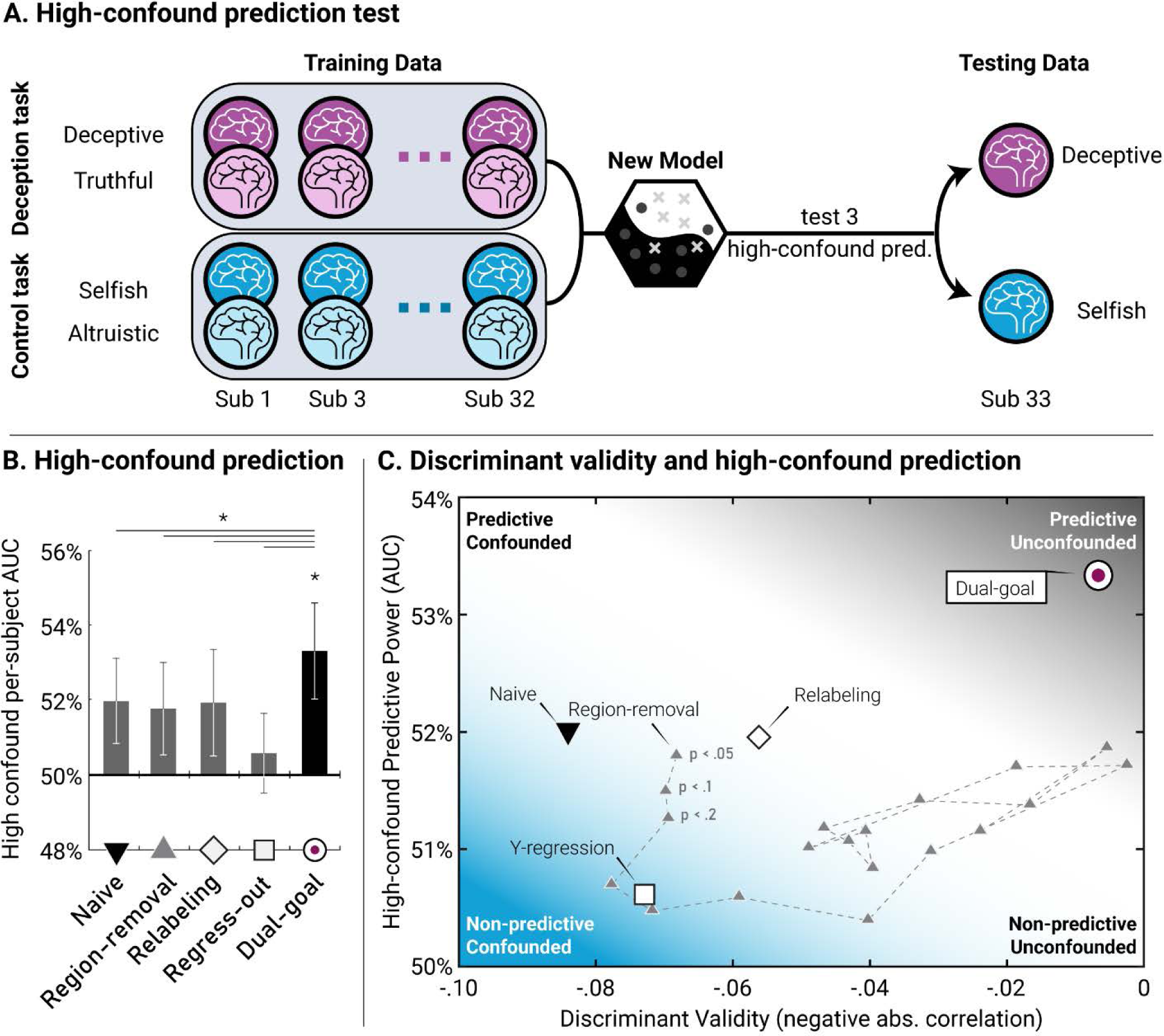
(**A**) Schematic of test 3: prediction in a high-confound test set in which lie trials are mixed with a confounded selfish but non-deceptive behavior. (**B**) Bar graphs of validity tests and high-confound prediction performances. Only dual-goal tuning shows significant discrimination between lie trials and selfish trials. Dual-goal tuning also showed significantly higher AUC than each of the other four methods at *p* < .05. (**C**) Discriminant validity (test 2) and high-confound prediction performances (test 3) of different neural predictors compared on a 2D plot. In contrast with dual-goal tuning, all other methods fail to adequately control nuisance correlation and hence result in low discriminability between highly confounded trials. * *p* < .05.

We found that, whereas dual-goal tuning was able to significantly distinguish between the two trial types (mean AUC = 53.3%, p = 0.0177), all other methods did not result in greater than chance rates of performance (**Figure 3B & C**). Furthermore, dual-goal tuning had significantly higher performance than each of the other four methods (*p* < .05 for all tests). Thus, consistent with the indirect evidence provided by our overgeneralization results above, these data provide positive evidence that dual-goal tuning was able to improve discriminant validity of the deception predictor.

### Neural systems underlying deception

Beyond prediction, the ability to detect and control for nuisance processes can also enrich our understanding of neural systems underlying deception. For example, previous meta-analyses of fMRI studies of deception have suggested the involvement of a network of regions in deception, including anterior insula, anterior cingulate, inferior frontal gyrus, inferior parietal lobule, and superior frontal gyrus (*21*, *41*, *42*). However, it is unclear the extent to which predictors based on these regions are vulnerable to the presence of confounding processes captured by our control task.

Using our deception task and searchlight MVPA (*43*), we corroborate previous meta-analytic findings (*21*) and show that regions such as dorsomedial prefrontal cortex (dmPFC) and precuneus contain signals that allow us to decode deception (**Figure 4A**). We used a searchlight with a radius of 2 voxels (33 voxels in a spherical ROI) and a partial least squares (PLS) algorithm with leave-one-subject-out cross-validation, followed by whole-brain permutation testing of significant predictive performance. We find evidence that overgeneralization at the ROI level is significantly more likely to occur than would be expected by chance, such that predictors trained on the deception task also generalize to the control task (permutation test *p* = .003; **Figure 4B**). Contingent on the stringency of the null criterion for meaningful overgeneralization, a more systematic examination of different cutoff values further shows that the percentage of voxels that overgeneralize ranges between 18 and 88 % (*p* < 0.05: 18%, *p* < 0.1: 28%, *p* < 0.2: 39%, *p* < 0.5: 66%, *p* < 0.8: 88%).

**Figure 4.**
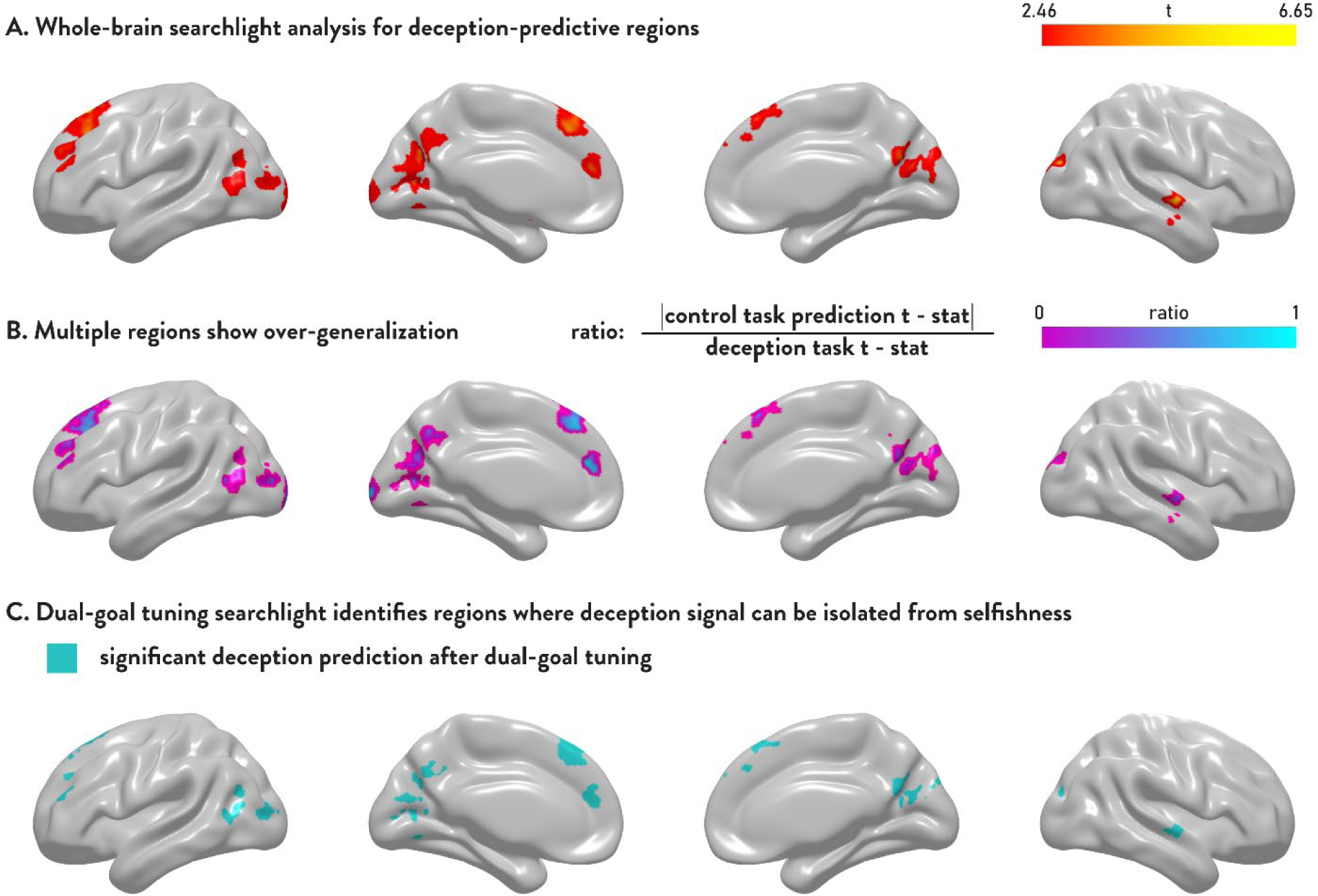
(**A**) Searchlight analysis for regions that can predict deceptive choices. Searchlight analysis with a radius of 2 voxels was performed across the entire brain to identify regions that significantly predict deceptive choices, as assessed by leave-one-out cross-validation. Regions with predictive performances significantly above 50% at the whole-brain correction level (permutation tested TFCE p <.05) are shown. (**B**) Cross-task generalization performance is measured for regions identified in **A**; the ratio of the t-test statistics is shown. Several regions that have high predictive power in panel **A** are also shown to have high generalization in panel **B**. (**C**) Dual goal tuning is applied at each searchlight to eliminate cross-task generalization and thereby identify regions that can significantly predict deceptive but not selfish choices (p<0.05).

Importantly, the extent to which the predictive regions were affected by post-training orthogonalization sheds light on the nature of neural signals in these regions. Some highly predictive regions, such as the left superior frontal gyrus and the left occipital pole, were no longer able to significantly predict deception after dual-goal tuning, suggesting that their predictive power was entirely driven by confounded signals (**Figure 4C**). In contrast, other highly predictive regions, such as the dmPFC, were able to retain predictive power even after orthogonalization, suggesting the co-mingling of distinct signals within these regions.

## Discussion

Deception is ubiquitous in nature. Male bowerbirds, for example, have been found to use optical illusions to enhance the attractiveness of their nests by arranging objects in a way that creates exaggerated size or depth perceptions (*44*). In humans, frauds such as investment scams cost consumers billions of dollars every year (*45*). Clinically, dishonesty is known to be an important characteristic of a number of mental and behavioral disorders (*46*) and has significant public health implications (*47*). However, despite its acknowledged importance, studies on the relationship between deception and the underlying biological mechanisms have long been discounted because of a historical lack of attention to its scientific foundations. As the National Research Council lamented in its 2003 report on the poor knowledge of diagnostic and psychometric properties of lie detection techniques, “More intensive efforts to develop the basic science in the 1920s would have produced a more favorable assessment in the 1950s; more intensive efforts in the 1950s would have produced a more favorable assessment in the 1980s; more intensive efforts in the 1980s would have produced a more favorable assessment now (*5*).”

Our work seeks to break this stalemate by providing such a scientific foundation. First, we build upon pioneering cognitive neuroscience studies in which participants could choose to lie, rather than being instructed to do so. By capturing the fact that honesty and deception are properties of the communicative signals that agents send to one another in the service of some economic or evolutionary benefit, meta-analyses of fMRI studies using these paradigms have identified a consistent set of brain regions in the lateral and medial PFC, as well as the anterior insula, that are more strongly engaged by deceptive compared with honest responses (*21*, *48*). More recent efforts have extended these findings by incorporating machine learning methods to predict behavior from brain activity, permitting researchers to avoid problems associated with reverse inference. Speer et al. (2020), for example, found that connectivity patterns in the self-referential thinking network were able to predict the honesty of participants during decision-making (*19*).

Building on these efforts, here we sought to explore a complementary aspect of prediction and generalization: the need to evaluate for the presence or absence of potentially confounding processes. Specifically, we leverage generalization tests in MVPA methods to test for the presence of confounding processes in a deception predictor. While generalization tests have become increasingly popular in cognitive neuroscience (*17*, *30*, *31*, *34*, *35*, *49*, *50*), to our knowledge it has not been applied in studies of deception. Our finding that signals underlying the neural predictor of deception are associated with non-deceptive, selfish choices supports long-standing validity concerns that predictive signals for deception can be driven by confounding processes.

In addition, our results suggest that using cross-task generalization to identify confounding signals can provide essential information about the construct validity in a manner that extends beyond within-dataset sensitivity and specificity (*17*, *21*). To clearly disambiguate these distinct contributions, we note that measures such as sensitivity or specificity address criterion validity using metrics such as the percentage of the total number of lies the predictor identifies, or the percentage of truths it falsely flags as lies. Cross-task generalization, on the other hand, probes the construct validity of the predictive signal by asking whether the predictive signal is correlated with unrelated measures, such as being significantly greater for certain types of truth trials (selfish ones) than other truth trials (altruistic ones) (*22*, *23*).

Our second contribution is to develop a novel approach to create predictors that do not demonstrate undesirable out-of-sample generalization when applied to a new task. While there have been studies in which an absence of generalization across two tasks has been used to show evidence for distinctiveness of two constructs (e.g., (*35*)), no methods to our knowledge have been developed that focus on eliminating an already existing generalization. Furthermore, while pioneering work on regression models and machine learning algorithms in neuroimaging has primarily addressed the goal of tuning model hyperparameters to improve predictive performance (*33*, *51*), for confounding signals overgeneralization requires reductions in performance, rather than improvements.

Our results suggest that controlling for overgeneralization can be achieved by addressing the predictor construction directly rather than altering what is included in the training data. Preprocessing the training data by removing the most confounded voxels, (i.e. region-removal) performs poorly when signals of interest and no-interest co-mingle at the voxel level. In such cases, region-removal can result in reduced power in predicting the variable of interest due to imperfect orthogonalization (**Figure 5A**). On the other hand, our dual-goal tuning procedure can be seen as a shearing transformation that reduces the inner product between the prediction map and the nuisance covariance (**Figure 5C**).

**Figure 5.**
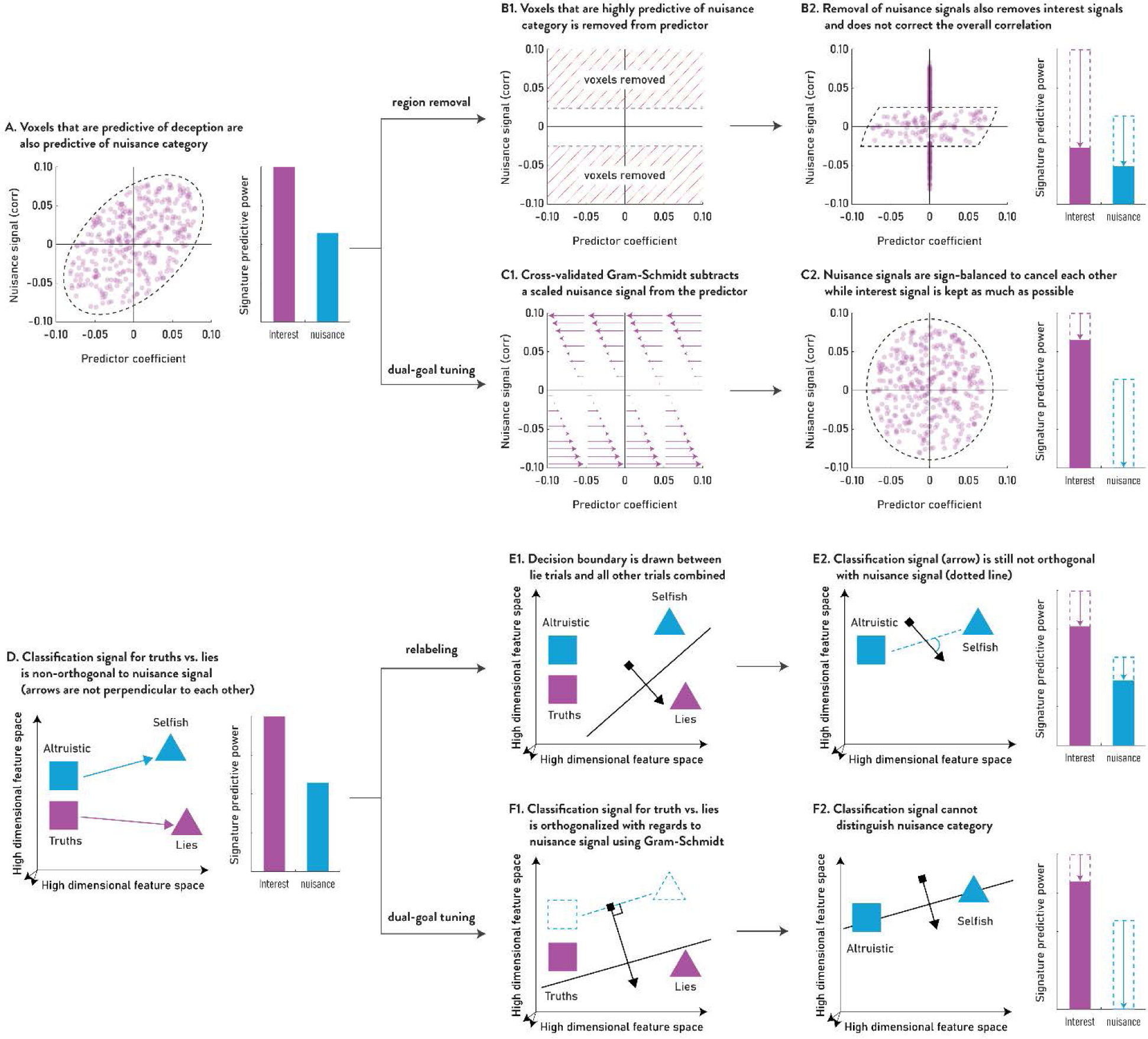
Comparison of region removal, relabeling, and dual-goal tuning in the presence of confounding processes. (**A**) Depiction of an underlying signal for which the signals of interest are confounded with nuisance signals. Voxels that carry highly positive signal for the signal of interest also carry highly positive signal for the nuisance signal, and vice versa for negative signals. (**B**) Depiction of the region-removal method, in which voxels that are strongly correlated with nuisance signals are removed before building a predictor. However, because the underlying non-orthogonality has not been solved, the region-removal approach is unlikely to achieve orthogonality. Furthermore, as the voxels that are correlated with nuisance are removed, so are the voxels that are correlated with signal of interest. (**C**) Depiction of the dual-goal tuning approach using Gram-Schmidt orthogonalization to correct the predictor. The shearing transformation is controlled by the orthogonalization hyperparameter so as to achieve zero out-of-sample predictive power for nuisance. (**D**) Depiction of a classification space in which the underlying signal of interest (purple arrow) is not orthogonal (perpendicular) to the nuisance signal (blue arrow). (**E**) Depiction of the relabeling approach, in which a best decision boundary is identified between the lie trials (purple triangle) and all other trials. While the decision boundary may be effective in labeling all truth trials as truths, the predictive signal (purple arrow) is still correlated with the nuisance signal such that selfish trials still receive higher prediction scores than altruistic trials. (**F**) Depiction of the dual-goal tuning approach, in which the predictive signal (purple arrow) is constructed under dual-goal tuning to be orthogonal (perpendicular) to the nuisance signal (blue dotted line) such that the predictor cannot reliably distinguish between altruistic and selfish trials.

In contrast, by introducing data from the control task to the training process without changing the optimization function, the relabeling approach attempts to find a decision boundary to distinguish the lie trials from all other trials. However, while this procedure enriches the examples of ‘truth’ trials in order to identify a better decision threshold, the predictive signal, which runs orthogonal to the decision boundary, is still correlated with the nuisance signal (**Figure 5E**). On the other hand, by conceptualizing the problem as a form of constrained optimization, dual-goal tuning orthogonalizes the predictive signal with regards to the nuisance signal such that the predictor cannot distinguish between the selfish and altruistic non-deceptive choices no matter where the decision boundary is placed.

Scientifically, we contribute by elucidating fundamental questions about the neural basis of honesty and deception. At the whole-brain level, the question is one of construct dissociability: can one adjust the signals of multiple regions in order to dissociate two related constructs? Our findings suggest that deception is indeed dissociable from other forms of social norm violation (*52–54*). Because activity in the preference condition can explain some, but not all, brain activity in the deception condition, these data highlight that understanding deception requires accounting for both self-interest and additional factors, such as the awareness of social norms and considerations relevant to their violation, that have dissociable neural representations.

Similarly, at a regional level, we address whether a signal from a given brain region is (i) specific to a certain construct, (ii) similar across constructs but dissociable, or (iii) common across constructs and non-dissociable. The searchlight analysis here reveals that regions such as the posterior temporal cortex and precuneus hold representations predictive of deception that are relatively unconfounded, while others, such as the medial PFC, hold presumptively multiplexed but dissociable representations that become evident when dual-goal tuning is applied. Importantly, yet other areas, including those in the occipital cortex and superior frontal gyrus, no longer predict deception after signals related to selfishness are removed. Thus, while overgeneralization is prevalent in regions implicated in previous meta-analytic studies of deception (*21*, *48*), at least some of these regions contain more specific signals.

More broadly, these data highlight the importance of discriminant validity in defining the cognitive processes underlying complex psychological phenomena (*15–17*, *55*). Currently, dissociating co-occurring or confounded processes requires an experiment that allows for orthogonal control of both processes, an approach that is not always possible. Failing that, studies have argued for the distinctiveness of mental processes by showing an absence of overgeneralization across datasets with naïve predictors (*34*, *35*). However, our results suggest that even when there is a considerable amount of overgeneralization, the underlying mental processes may be distinguishable. Our methodology may therefore be useful in dissociating common co-occurring processes, especially if the orthogonality must be established post-hoc or used as a complement to specific task designs, such as cases involving working memory and attention (*56*), valuation and salience (*57*), or valence, arousal, and emotions (*49*). In computational psychiatry, our approach may aid in constructing neural biomarkers specific to a particular diagnosis without relying on covariates of no interest (*58*, *59*).

The ability to test and control for the presence of confounding processes also opens the door to a number of new avenues of research. First, additional work is needed to account for individual-level heterogeneity in out-of-sample predictions. While in principle it is possible to build separate predictors, it may well be infeasible to obtain training data for every individual, particularly for practical applications such as those in forensic and mental health settings. Depending on the nature of the questions, one can either treat individual heterogeneity as a factor of no interest in order to identify core components of deception, or more fully characterize the neural heterogeneity of deception in order to account for it when making out-of-sample predictions.

Finally, our findings point to the value of future work that can further decompose underlying neurocognitive mechanisms of deception (*13*, *25*, *26*, *48*, *60*). In particular, inclusion of additional control tasks with more basic tasks focusing on arousal, reward processing, or belief inference processes can yield additional valuable insight by decomposing the underlying mechanisms associated with deception (*4*, *21*). That is, in addition to testing discriminant validity with our control task, we can gain further knowledge about convergent validity using tasks that target specific cognitive processes (*22*). Such a strategy can also be used to elucidate mechanisms underlying forms of deception that we did not address in the current study, for example altruistic deception or deception through omission. A comparison of these different forms of deception can in turn elucidate common and distinct underlying mechanisms. Thus, despite important limitations, our findings represent a meaningful advance given the significance of the questions and the protracted nature of the challenges involved (*5*).

## Acknowledgements

This study was supported by grant funding from the National Institute of Mental Health (R01 MH112775 to MH and ASK), the National Institute on Alcohol Abuse & Alcoholism (R01 AA026587 to ASK and MH), the National Science Foundation (1851902 to MH), STI2030-Major Projects (2022ZD0205100 to LZ), and NSFC (32325023 and 32071095 to LZ). The authors declare no competing interests.

## Supplemental Materials

### Methods

#### Participants

Forty people provided informed consent and participated in the experiment (16 males; age = 20.8 years ± 2.6 years (mean ± sd)). Seven participants were excluded from data analysis because they exhibited extremely one-sided choices in the task and therefore lacked sufficient behavioral variation to predict (see below). As a result, 33 participants were included in final data analysis. All experimental procedures were approved by the Institutional Review Board of Peking University.

#### Experimental Design

We used a messaging task in which participants, in the role of the sender, chose one of two messages to send to a recipient (*1*, *2*). On each trial, participants saw two monetary allocations on the screen (A and B). One of the monetary allocations provided more money for the participant than the recipient, while the other allocation provided more money for the recipient than the participant. In the main deception task, participants could choose between two messages: ‘option A will make you more money than option B’ and ‘option B will make you more money than option A’. Given the allocations A and B, only one of these two messages was true; the other message was deceptive but yielded more money for the participant. Hence, participants made decisions between sending an altruistic honest message and a deceptive selfish message. In the control task, while the monetary allocations were the same, the message that participants could send was either ‘I prefer that you choose option A’ or ‘I prefer that you choose option B’. In this case, neither message was deceptive; instead, each was a statement of preference. All participants completed 40 trials for each task (see **supplemental Table S1** for the allocations presented on individual trials).

Significantly, for both conditions, participants were aware that as the sender, they could see the monetary options but could not make the choice between them, while the receiver could not see the monetary options but was responsible for the choice between them. This important manipulation renders the receiver completely reliant upon the sender for any information about the choice, preventing the receiver from using payoff information to infer the sender’s behavior. In addition, senders were aware both that in the deception condition, recipients would never know the truth value of the message they received, and that they as the sender were paired with a separate receiver on each trial, limiting influences of any one trial on subsequent trials (*1*, *2*).

#### Image acquisition, preprocessing

MRI images were collected with a Siemens 3T Prisma scanner with a 32-channel head coil. High-resolution T1-weighted anatomical images were acquired using an MPRAGE sequence [repetition time (TR) = 2,530ms; echo time (TE) = 2.98 ms, 512 axial slices, 1 x 0.5 x 0.5mm voxels; 192 x 448 matrix). T2*-weighted functional images were acquired using an EPI sequence with 32 axial slices 3 x 3 x 4.4mm voxels, 64 x 64 matrix, TR = 2,000 ms, TE = 30 ms. All images were preprocessed using fMRIPrep 20.2.1 (*3*). EPI sequences were skull-stripped, co-registered using boundary-based registration with nine degrees of freedom, head-motion corrected via six degrees of freedom, slice-time corrected, and normalized to a 4mm MNI space. Before running the activity estimation procedure below, the images were smoothed with a FWHM 8mm Gaussian kernel.

#### Neural activity estimation for decoding analyses

Neural prediction was performed at two levels. At the subject-level, activity for trials of the same category was estimated together, such that one activity image per category was estimated for each run. At the trial-level, each trial’s activity was estimated separately. The former approach, compared to the latter, enjoys the benefits of higher SNR, as activity is averaged over several trials and hence is more stably estimated. However, run-level estimation uses knowledge of trial groupings. Although the researcher need not know the identities of the groupings, in practice relatively few situations may be amenable to group-category predictions when researchers do not know how each trial should be grouped.

For subject-level predictions, we used a GLM with one regressor for all trials in which the participant made truthful/altruistic choices, and one regressor for all trials in which participants made deceptive/selfish choices. For single-trial predictions, the GLM consisted of a separate regressor for each trial (beta-series regression (*4*)). Regressors were modeled with an impulse function time-locked to the button press and convolved with a double-gamma HRF to simulate BOLD (blood-oxygen level dependent) activity. Nuisance regressors included the top 10 PCA components derived from the combined CSF + white-matter masks (a_comp_cor, provided by fMRIPrep), as well as 24 motion parameters (6 affine transformations for each TR and the previous TR, and their squares (*5*)).

#### Leave-one-out cross-validation

All prediction exercises in this paper used leave-one-subject-out cross-validation. That is, a predictor was trained on the 32 participants’ main deception task, and the trained predictor was used to predict the choices of the left-out participants’ main deception task and the control task. This procedure was repeated 33 times until all participants’ data had been predicted. This leave-one-out procedure allowed us to perform cross-validation while retaining as much training data as possible, at the cost of computation time from repeating the prediction 33 times.

Within the training data (i.e., 32 participants), we trained whole-brain neural predictors using the Thresholded Partial Least Squares (T-PLS) algorithm (*6*). T-PLS was chosen because it has been shown to be computationally fast, yet highly powerful in whole-brain predictions. T-PLS employs two hyperparameters that require tuning for optimal predictions: the number of PLS components, and the thresholding level. The number of PLS components generally controls the regularization, as using fewer components leads to greater dimension reduction in the data. The thresholding level determines variable selection, as it controls how many of the non-predictive voxels will be removed from the predictor. Based on factors including the number of trials and the size of the searchlight, we explored a number of components from 1 to 5, and a thresholding level from 50% to 100%. In order to tune the hyperparameters of T-PLS, we employed another leave-one-out cross-validation within the training data (i.e., nested cross-validation). Specifically, we trained a T-PLS model on 31 participants and used it to predict the left-out participant in the training data while varying the hyperparameters.

#### Regress-out methods

Regression is widely used to orthogonalize two vectors. In the current study, we have two datasets {***X****_1_*, ***Y****_1_*}, and {***X****_2_*, ***Y****_2_*}, where ***X*** is a matrix of brain images (with rows comprised of trials and columns consisting of voxels) and ***Y*** is the behavior (a single column vector of binary choices). In this situation, one could potentially use regression in multiple ways. However, it is important to first note that, from a practical point of view, all regress-out approaches listed here are less versatile than other methods discussed in the manuscript. The regress-out methods require two datasets to be of the exact same size and to have a coupled relationship between observations. In other words, all observations of ***Y****_2_* need to be matched one-to-one with all observations of ***Y****_1_* in order for the regression to work. In the case of the current study, because the two tasks are isomorphic and have trials with the same payoff structure, a reasonable pairing can be constructed to enable trial-level prediction. However, if one considers a more general study in which one wishes to rule out confounding signals generated by other tasks (e.g. a stroop task, a delay discounting task, or others), it becomes impossible to identify which trials of the two unrelated tasks best correspond to each other.

*The ‘Y-Y regress-out’.* Perhaps the most intuitive of all regress-out methods is to regress behavior ***Y****_2_* (i.e. selfishness) out of behavior ***Y****_1_* (i.e., deception), with the hopes that removing the behavioral co-variation also results in neurally orthogonal predictors. Conceptually, the ‘Y-Y regress-out’ approach assumes that the *only* reason for over-generalization is the similarity in behavior between the ***Y***s, and that the underlying neural processes (i.e. the ***X***s) are independent. As shown in the main manuscript, this regress-out method did not successfully remove overgeneralization, suggesting that the underlying neural representations include shared components. As a formal example for why this method might frequently fail, one could conceive of two uncorrelated behaviors ***Y****_1_* and ***Y****_2_* with a corresponding neural signal ***X*** that reflects a sum of ***Y****_1_* and ***Y****_2_*. In such a case, regardless of the orthogonality of ***Y****_1_* and ***Y****_2_*, a predictor based on ***X*** will predict both ***Y****_1_* and ***Y****_2_*.

The failure of this method in the current manuscript also has implications for creating orthogonal neural predictors (i.e., predictors that don’t over-generalize) through factorial experimental designs. For example, Lee et al. (*7*) used a two-by-two factorial experimental design in which participants were asked to imagine scenarios that were either vivid/non-vivid with positive/negative valence. From this balanced design, they were able to create a neural predictor of imagination vividness that did not predict valence, as well as a neural predictor of imagination valence that did not predict vividness. While the orthogonality of the ***Y***s (vividness and valence) was sufficient to result in orthogonal predictors in their case, it was also possible that orthogonality could not be achieved had there been a number of brain regions whose signal was influenced by both vividness and valence.

*The ‘X-Y regress-out’.* Another possible approach is to remove from ***X****_1_* (i.e., deception brain data) any correlation with ***Y****_2_* (i.e., selfishness behavior). This choice is motivated by the fact that if all variations related to selfishness are removed from neural data, no linear combination of neural data can produce an outcome that is correlated with selfishness. This orthogonality, however, relies on being able to provide only pre-cleaned neural data, which is not tenable in out-of-sample prediction. If a predictor is trained on brain data from which the selfish signal has been removed, it will pick up the most predictive voxels without knowing that many of those voxels also contained nuisance signals prior to regress-out. Hence, when the time comes to use the predictor in a real test – i.e. data that has not been pre-cleaned – the predictor will utilize voxels that also identify confounding processes.

*The ‘X-X regress-out’.* The last approach is to remove the neural variance in ***X****_2_* (control task) from ***X****_1_* (deception task). This method is similar to the aforementioned ‘X-Y regress-out’ except that instead of regressing out ***Y*** from all voxels of ***X***, each voxel of ***X****_2_* would be regressed out from their corresponding voxel in ***X****_1_*. The difficulty with this method is the same as that in ‘X-Y regress-out’: the out-of-sample data does not come cleaned.

#### Overgeneralization and dual-goal tuning

Dual-goal tuning can be considered as a generalization of a constrained optimization problem since the goal is to optimize the deception predictive performance while satisfying the constraint that the predictor map has zero inner product with the nuisance task covariance. The novel aspect of this method comes from having to satisfy the constraint for out-of-sample, rather than in-sample, prediction. We found that in-sample orthogonalization of the predictor through constrained optimization always leads to some remaining positive cross-task generalization (see below). In the machine-learning literature, a number of studies have sought to improve the validity of prediction models by similarly constraining the kinds of signals used for prediction. Direct application of such methods to our problem was limited, however, as those methods explicitly require the provision of a guiding cost function for ‘correct’ signals (*8*, *9*). In our case, the correct signal – i.e. the neural signature of deception – is unknown in its form or loci. Instead, our attempt to reduce the overgeneralization can be thought of as a negative guiding cost function that penalizes what ‘incorrect’ signals do (i.e., overgeneralize).

As stated in the main manuscript, given two datasets (***X****_1_*, ***Y****_1_*) and (***X****_2_*, ***Y****_2_*), cross-task generalization occurs due to the similarity between the predictor map b and the nuisance covariance map ***C****_2_* = ***X****_2_^T^**Y**_2_*. Therefore, an intuitive solution to suppress such generalization is to add a term to the cost function that penalizes the inner product between the two maps. In the case of ridge regression, for example, the modified loss function would be as follows:

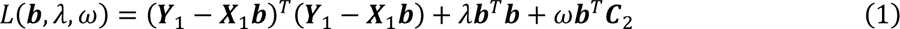

The hyperparameter *λ* controls the shrinkage of the coefficients and the hyperparameter *ω* controls cross-task generalization. In normal ridge regression, the *λ* that maximizes cross-validation performance is chosen. In this case, the set of hyperparameters (*λ*, *ω*) should be chosen to maximize prediction performance while cross-task generalization is held at null. While elegant in form, such a formulation of the cost function has two shortcomings. Firstly, it limits the dual-goal tuning approach to cost-function based methods and excludes data-reduction approaches such as partial least squares or principal component regression. Secondly, the computational burden can be significant as the dual-goal tuning process expands the hyperparameter space to be searched. The second problem is especially acute for *ω*, as it needs to be precisely tuned for each of the *λ* considered. For these two reasons, we consider a more generalized two-step approach below.

To motivate our two-step approach, we derive the analytical solution for the ridge cost function above:

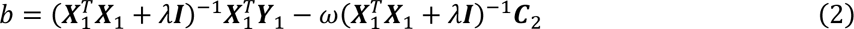

This solution can be understood as the sum of a naïve ridge predictor 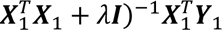 and a correction term 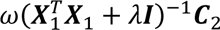. From this formulation, we consider a more explicit approach in which we start with a naïve predictor, then apply an orthogonalizing correction post-hoc. Here we use a Gram-Schmidt approach in which the map b is first constructed naïvely, and then orthogonalized with regards to ***C****_2_*:

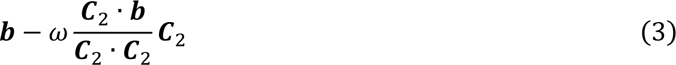

The Gram-Schmidt procedure subtracts a scaled version (i.e., projection) of the vector ***C****_2_* from vector ***b*** and makes it orthogonal to ***C****_2_*. With this formulation, the initial predictor creation step is separated from the orthogonalization step, enabling any prediction algorithm to be used for the initial predictor.

Importantly, while *ω* is typically equal to 1 for in-sample orthogonalization via Gram-Schmidt, in our case the orthogonalization will be performed using the training sample’s estimate of ***C****_2_*, which will be different from the out-of-sample’s true ***C****_2_*. To demonstrate the necessity for the scaling factor *ω*, let the in-sample’s estimate of ***C****_2_* be denoted as 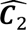. Then, if we orthogonalize our predictor with regards to 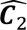, the out-of-sample dot product between our orthogonalized predictor and true ***C****_2_* is as follows:

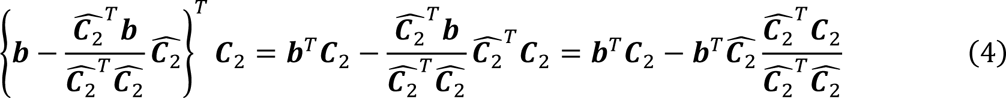

In the last part of the equation, ***b***^*T*^***C***_2_ and ***b***^*T*^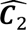 should be equal to each other in expectation, but the ratio of 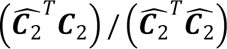 will be less than 1 in expectation given that the denominator is a dot product of the same vector while the numerator is that of different vectors of roughly similar norms (Cauchy-Schwarz inequality). Hence, the orthogonalization will likely be insufficient, which means that the hyperparameter *ω* should be greater than 1 to compensate for this effect.

### Whole-brain predictor before and after dual-goal tuning

The final whole-brain predictor of deception, trained from all data using T-PLS and dual-goal tuning, shows interpretably clustered coefficients across the entire brain (**Figure S1**). Strongly positive-weighted regions include the dorsomedial PFC, the dorsolateral PFC, and ventromedial PFC, while the precuneus was notably negative-weighted. These regions correspond well with the areas identified in the searchlight analysis (Figure 4). Notably, orthogonalizing the naive neural predictor (**Figure S1**, panel A) by subtracting the weighted covariance map from the control task (**Figure S1,** panel B) does not substantially change the predictor’s final appearance (**Figure S1**, panel C). Even as it implements the Gram-Schmidt orthogonalization, the shearing transformation maintains the general pattern of coefficients, resulting in a final map that retains the interpretability of the naive map.

**Figure S1.**
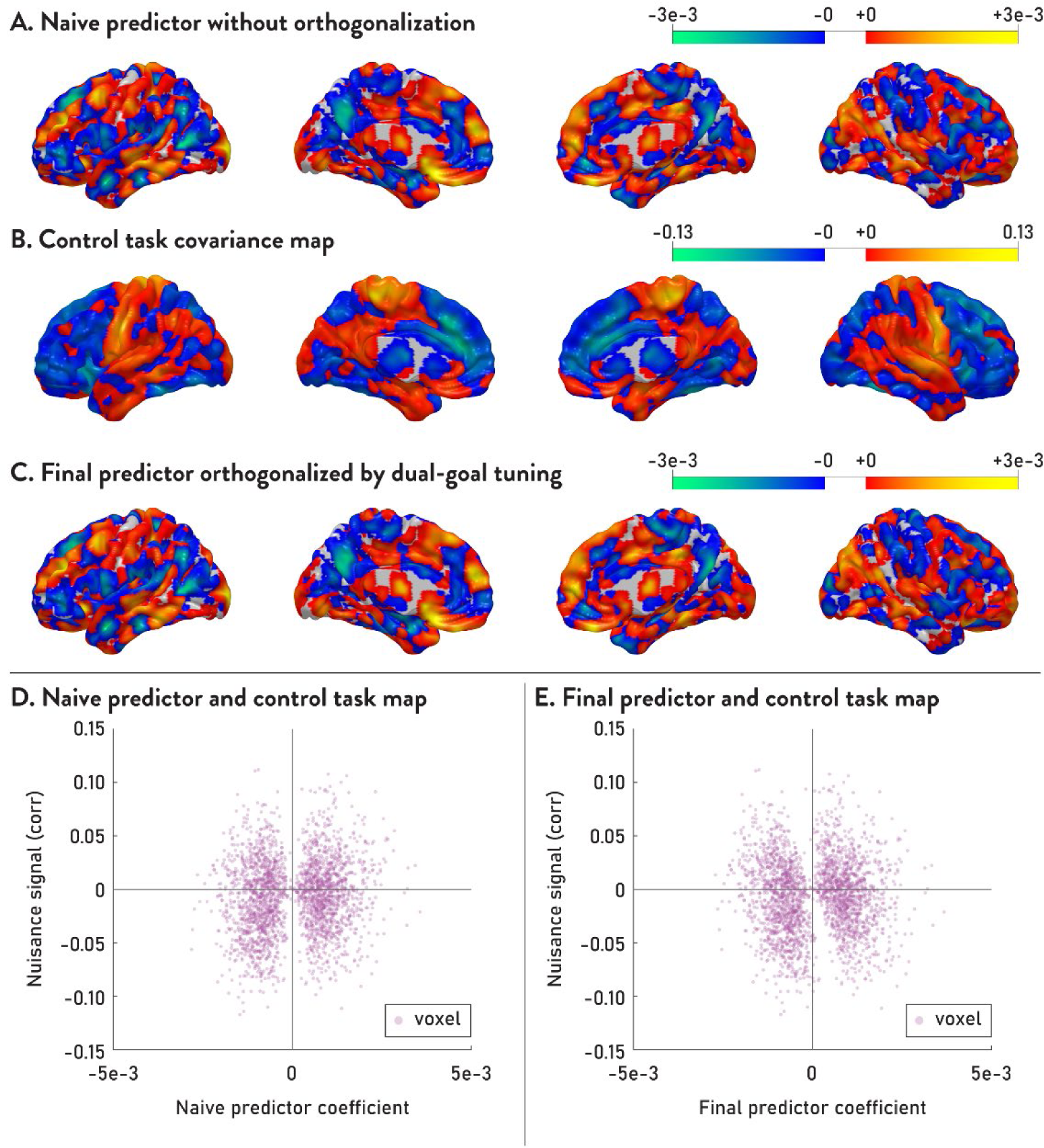
Final whole-brain predictor before and after dual-goal tuning. Panel A shows the naive whole-brain predictor created using the T-PLS algorithm on all 33 participants’ data, for visualization purposes. (Note that for the cross-validation analyses described elsewhere, the predictor was constructed using a leave-one-out procedure that included 33 different sets of 32 training subjects and one test subject.) The dual-goal tuning approach subtracts a scaled version of the covariance map derived from the control task (panel B) from the naive map (panel A). The resulting map, shown in panel C, has an overall appearance quite similar to the naive map in panel A. To explain this similarity, panels D and E show, for the naive predictor and the final predictor respectively, the scatterplots of the predictor coefficients (abscissa) and nuisance signal correlation (ordinate) for every voxel in the brain. As reflected in the apparent rotation of the data from panel D to panel E, dual-goal tuning implements a shearing transformation that reduces the size of the predictor coefficients for voxels in the first and the third quadrants, while increasing them for voxels in the second and fourth quadrants, thereby achieving orthogonality as defined by the dot product. The fact that the general pattern of coefficients remains largely unaltered results in visually negligible differences despite a significant change in generalization properties.

### Subject-level prediction and validity test performances of various methods

As in the trial-level prediction results shown in the main manuscript (Figure 2G & Figure 3B), all methods used retained significant predictive performances at the subject-level for the within-task prediction (test 1), but only dual-goal tuning resulted in meaningful elimination of cross-task generalization (test 2). As a result, only dual-goal tuning was able to successfully distinguish lie trials from selfish trials in high-confound prediction (test 3). Regress-out methods could not be included in subject-level prediction as the coupling relationship cannot be established between the observations of two tasks when the observations are grouped by the participants’ choices.

**Figure S2.**
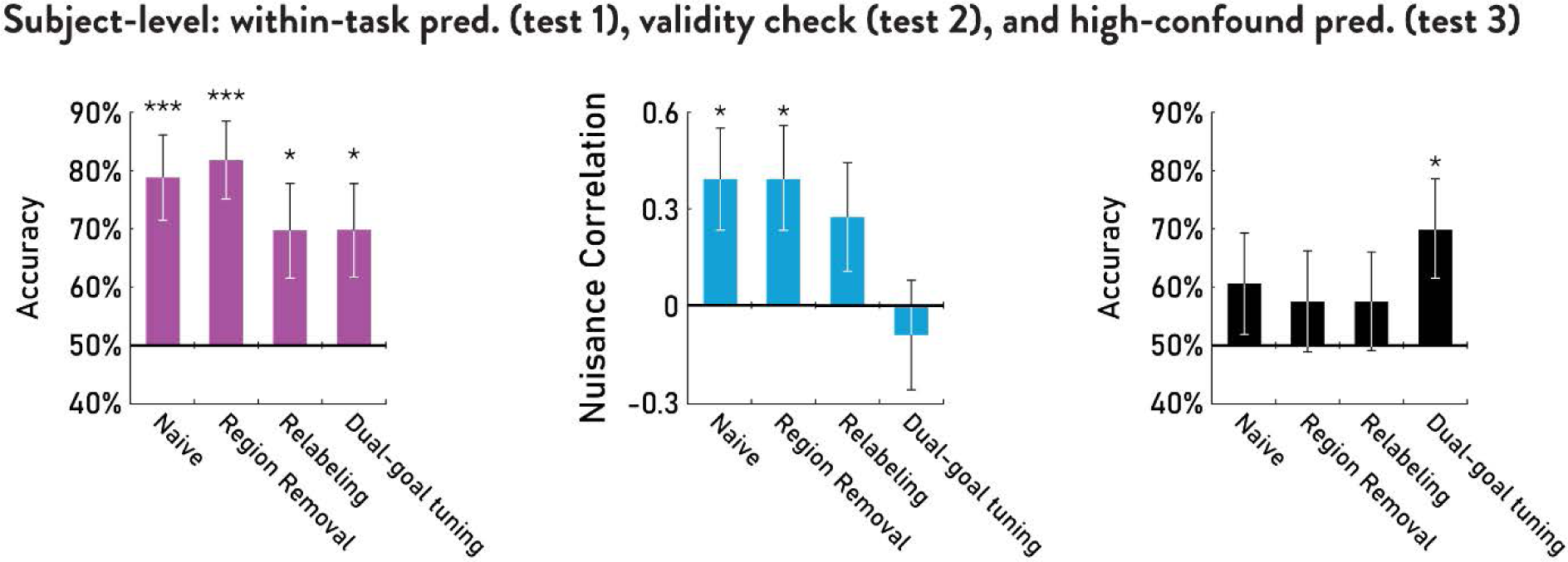
Bar Graphs of subject-level tests for within-task prediction (test 1, left in purple), cross-task validity check (test 2, middle in blue), and high-confound prediction (test 3, right in black). * *p* < .05, *** *p* < .001.

### Monetary allocations on experimental trials

**Table S1.** Monetary allocations on experimental trials: each subject was presented with the following 46 payoffs twice, once in the deception task, and once in the control task. In 40 out of 46 payoffs, one of the two options was more beneficial to the participant (sender) while the other option was more beneficial to the counterpart (recipient). In 6 out of 46 payoffs, one of the options was beneficial to both sender and recipient. These trials were included as a quality check, as the participants should choose the mutually beneficial option.

| Option A |  | Option B |  |
| --- | --- | --- | --- |
| Self | Other | Self | Other |
| 25 | 16 | 10 | 19 |
| 10 | 10 | 5 | 15 |
| 15 | 25 | 25 | 15 |
| 20 | 20 | 5 | 35 |
| 10 | 25 | 25 | 10 |
| 10 | 21 | 30 | 19 |
| 10 | 10 | 15 | 5 |
| 10 | 25 | 30 | 15 |
| 19 | 10 | 16 | 25 |
| 14 | 10 | 15 | 11 |
| 12 | 8 | 6 | 11 |
| 15 | 10 | 20 | 9.99 |
| 5.99 | 10 | 6 | 5 |
| 15 | 25 | 16 | 5 |
| 5 | 30 | 30 | 5 |
| 19 | 30 | 21 | 10 |
| 20 | 5 | 15 | 30 |
| 21 | 10 | 20 | 30 |
| 35 | 5 | 20 | 20 |
| 10 | 5.99 | 5 | 6 |
| 8 | 5 | 7 | 10 |
| 8 | 5 | 5 | 20 |
| 15 | 25 | 14.99 | 30 |
| 11 | 5 | 10 | 15 |
| 25 | 15 | 30 | 14.99 |
| 15 | 20 | 10 | 5 |
| 9.99 | 20 | 10 | 15 |
| 10 | 7 | 5 | 8 |
| 10 | 21 | 30 | 20 |
| 5 | 30 | 25 | 28 |
| 25 | 15 | 11 | 22 |
| 15 | 25 | 22 | 11 |
| 19 | 35 | 22 | 5 |
| 35 | 19 | 5 | 22 |
| 10 | 21 | 11 | 1 |
| 25 | 15 | 5 | 16 |
| 12 | 8 | 11 | 6 |
| 20 | 10 | 30 | 15 |
| 20 | 5 | 5 | 8 |
| 1 | 11 | 21 | 10 |
| 30 | 15 | 5 | 20 |
| 5 | 6 | 8 | 10 |
| 25 | 10 | 15 | 30 |
| 15 | 10 | 5 | 11 |
| 28 | 25 | 30 | 5 |
| 20 | 20 | 10 | 15 |

